# Defoliation-induced compensatory transpiration is compromised in *SUT4*-RNAi *Populus*

**DOI:** 10.1101/2020.01.13.905406

**Authors:** Scott A. Harding, Christopher J. Frost, Chung-Jui Tsai

## Abstract

Leaf sucrose contents are high in species of the genus *Populus* and other temperate tree taxa. Sucrose is subcellularly compartmentalized, but assumptions about the physiological relevance of such partitioning remain largely unexplored. In this study, the effects of partial defoliation treatments on water uptake, leaf gas exchange properties, non-structural carbohydrate abundance in source and sink organs, and growth were compared in poplars with normal or impaired intracellular sucrose trafficking. The tonoplast sucrose transporter PtaSUT4 is well expressed in leaves of *P. tremula* × *P. alba* (INRA 717-IB4), and its inhibition by RNA-interference (RNAi) is known to affect leaf sucrose abundance. After partial defoliation, maximum photosynthesis rates increased while intercellular CO_2_ decreased with trajectories that were similar in wild type and *SUT4-*RNAi lines. Leaf transpiration increased more robustly in wild-type than RNAi plants, while leaf water content increased more in RNAi lines. Stomatal conductance did not differ between genotypes, nor did it increase with defoliation. Post-defoliation reductions in steady-state levels of sucrose, the major hexoses (glucose and fructose) and starch were similar in wild-type and *SUT4-*RNAi shoot sinks. Water uptake and stem growth after partial defoliation were not as well sustained in RNAi as in wild-type plants. The data suggest that vacuolar efflux of sucrose by PtaSUT4 more clearly facilitated adjustments in water uptake than sucrose export following leaf removal.

## Introduction

Sucrose and its hexose breakdown products contribute significantly to the high leaf osmolalities often found in temperate tree species like *Populus* (Rennie and Turgeon 2009; Slewinski et al. 2013). High osmolality is thought to enable turgor maintenance under conditions of low plant hydraulic conductivity while obviating a need for energy-dependent active phloem loading of sucrose in these species that depend on passive diffusion of sucrose for long distance transport (Davidson et al. 2011; Fu et al. 2011). Mesophyll sucrose resides in both the vacuole and the cytosol but little is known about how sucrose trafficking across the tonoplast might impact osmotic regulation. In addition to leaf turgor maintenance, there is the question of how sucrose partitioning might impact sink carbohydrate provisioning. Tonoplast trafficking of sucrose has long been considered important for the maintenance of cytosolic sucrose at high enough concentrations to support passive diffusion into the phloem (Turgeon and Medville 1998). This would presumably mean that sucrose accumulating during the day and stored in the vacuole would be released at night in order to sustain high cytosolic levels and diffusion to the phloem for sink provisioning. Interestingly, it is starch rather than sucrose levels that become depleted at night in *Populus* (Zhang et al. 2014). From these observations, it is unclear whether tonoplast sucrose trafficking in *Populus* is as important for export to sinks as it may be for plant hydrodynamics such as transpiration.

Sucrose compartmentalization involves its uptake into the vacuole by the tonoplast monosaccharide proton antiporter and its efflux into the cytosol by the tonoplast sucrose proton symporter SUT4 (Reinders et al. 2008; Schulz et al. 2011; Schneider et al. 2012). In *Arabidopsis* where phloem loading is active and mediated exclusively by plasma membrane SUTs (Ayre 2011; Liesche 2017), *AtSUT4* is very weakly expressed, representing less than 5% of leaf *SUT* transcript abundance (Lloyd and Zakhleniuk 2004; Schneider et al. 2012). In the monocot sorghum, the tonoplast SUT4 is as well expressed as plasma membrane SUTs in source leaves and thought to facilitate active phloem loading (Milne et al. 2013). Leaf sucrose exporting ability and plant growth are impaired in rice mutants defective in the SUT4 ortholog *OsSUT2*, which led to the conclusion that passive transport is important in rice and that OsSUT2 drives sucrose export to the sink organs (Eom et al. 2011; Eom et al. 2012). In the poplar hybrid *P. tremula* × *P. alba* INRA 717-1B4 where sucrose is not actively loaded into the phloem (Fu et al. 2011; Zhang et al. 2014), *PtaSUT4* is the most highly expressed SUT in leaves (Payyavula et al. 2011). RNAi inhibition of *PtaSUT4* led to elevated leaf sucrose levels consistent with vacuolar sequestration, but unlike the case for the *ossut2* mutant of rice, plant growth was not impaired (Payyavula et al. 2011). Besides the possible role in sucrose export to sinks, the potential for SUT4 to modulate transpiration has not been explored.

*AtSUT4* expression is more readily detected in roots than shoots, and evidence for osmotic responsiveness of the *AtSUT4* promoter independent of sucrose has been described (Schneider et al. 2012). Increased sucrose levels in *Arabidopsis* roots following osmotic stress or ABA treatment have been attributed to shoot AtSUT4 in conjunction with plasma membrane phloem-loading SUTs (Gong et al. 2015). However, substantial stress-induced root sucrose increases were also noted in null mutants of those transporters (Gong et al. 2015). When *Arabidopsis* plants were grown from seed under mild water deficit conditions, sugar levels increased in shoots and roots, but *AtSUT4* was not among the *SUTs* that exhibited transcript level increases (Durand et al. 2016). This contrasts with large *PtaSUT4* expression decreases that occur in poplar plants grown under chronic, mild water deficit conditions (Frost et al. 2012). Consistent with the notion of functional versatility in *Populus, PtaSUT4* expression has been found to respond differently to chronic and acute water stress (Frost et al. 2012; Xue et al. 2016; Pagliarani et al. 2019). In addition, aquaporin expression differs between drought-stressed wild-type (WT) and *SUT4*-RNAi poplars, consistent with a link between sucrose trafficking and changes in water flux (Xue et al. 2016). Together, it appears that the physiological role of SUT4 includes export of sucrose to distant sinks for their nourishment, modulation of hydraulic properties that affect transpiration, or a delicate balancing of both.

As large, long-lived perennials, broadleaf tree species constantly undergo changes in source:sink ratio which, like stresses, trigger compensatory changes in photosynthesis, water uptake and carbohydrate utilization as plants adjust (Turnbull et al. 2007; Wiley et al. 2013). Growth reductions and mortality can follow repeated defoliations, but photosynthesis and transpiration can increase on a leaf area or per tree basis following partial defoliation or stand thinning (Breda et al. 1995; Eyles et al. 2013). In several studies, the transpiration increases were accompanied by more negative leaf water potentials and stable leaf hydraulic conductivity although the relationships vary across species and with stress severity (Brodribb and Holbrook 2006; Eyles et al. 2013; Liu et al. 2014). Diurnal adjustments in leaf water potential and hydraulic conductivity as transpiration oscillates, even under well-watered conditions have also been reported (Simonin et al. 2015). Variation among poplar genotypes with respect to leaf water potential adjustments and growth under stress conditions has been described, and probably depends on multiple underlying controls (Tschaplinski et al. 2006). The present work utilized pot-grown WT and *SUT4*-RNAi saplings to assess whether SUT4-mediated sucrose efflux from the vacuole potentially contributes to compensatory mechanisms deployed in tree leaves following defoliation.

## Materials and Methods

### Plant propagation and growth

Generation and characterization of the *SUT4*-RNAi lines in the hybrid poplar *P. tremula* × *P. alba* INRA 717-1B4 background (717) were previously described (Payyavula et al. 2011; Xue et al. 2016). Single-node cuttings were grown in a glasshouse in 4 gallon tree pots (Hummert International) containing commercial soil mixture (Fafard 3B) supplemented with Osmocote (15-9-12 NPK 4-month release) as previously described (Frost et al. 2012; Xue et al. 2016). Two experiments were conducted, the first, an 8-day experiment during May and June 2012; the second, a 16-day experiment during May and June 2016. The second experiment was necessitated because samples from the first experiment became degraded due to freezer malfunction before starch, condensed tannin and phenolic glycosides were measured. Outdoor conditions were essentially identical for both experiments. In both cases, evaporative cooling pads kept daytime temperatures below 35°C at canopy height. Plants were watered daily throughout the growth and treatment phases of the experiments.

### First defoliation experiment

For the first experiment, sixteen vegetatively propagated copies of WT and two RNAi lines (F and G) of roughly uniform height (∼1.5 m) were randomly assigned to three defoliation treatments and a control treatment with no defoliation (N=4 plants per genotype x treatment combination). At the time of defoliation, upper stems were marked at the position of leaf plastochron index zero (LPI-0) according to established developmental index criteria for *Populus* (Larson and Isebrands 1971). For the 25% and 50% partial defoliation treatments, respectively, every fourth or every other leaf and its associated bud scale below LPI-0 was removed. For the 100% defoliation treatment, all leaves below LPI-0 were removed. Experimental plants were maintained with normal watering for 8 days at which time photosynthetic parameters were measured and tissue harvesting was carried out as detailed below.

### Photosynthesis

Leaf photosynthesis, stomatal conductance and transpiration of WT and *SUT4*-RNAi transgenic plants were determined at midday and early afternoon using a Licor LI-6400XS (LiCor, Lincoln, NE) as described previously (Frost et al. 2012). LPI-6, a newly expanded source leaf, was used for the measurements. Photosynthesis, stomatal conductance and transpiration rates were determined at a single, saturating light intensity of 1500 μmol/m2/s.

### Biomass analysis and tissue sampling

Each plant comprised a single unbranched stem. No syleptic bud release along the stem occurred during the treatment phase. Immediately prior to harvest, a fully expanded source leaf at LPI-10 was weighed and snap frozen in liquid nitrogen for metabolite analysis. A stem section between LPI-15 and 20 was debarked after weighing, and both bark and wood fractions snap frozen for metabolite analysis. Upon harvest, fresh weights of the remaining leaves and stems were obtained. Tissues were then placed in a forced-air oven and dried at 100°C for 48 hr for dry weight determinations.

### Sucrose and hexose analysis

Snap-frozen leaves were lyophilized for 48 hr (FreeZone 2.5, Labconco), then ground through a 40 mesh sieve using a Wiley Mill (Thomas Scientific). Aliquots of the coarse powder were further ball-milled in a Mini Bead-beater (Biospec 3110Bx) at intensity setting 25 for two cycles. Ten mg of the lyophilized powder was suspended in a microtube with 500 µl methanol:chloroform (1:1, v/v) containing adonitol as internal standard, and sonicated for 15 min in a sonic bath with pre-chilled water (4°C). Deionized water (200 μl) was then added to the tubes and samples vortexed and re-sonicated for 5 min. After centrifugation, 10 μl of the upper aqueous-methanol phase was evaporated to dryness in 200 μl glass microserts, and derivitized for Gas Chromatography-Mass Spectrometry (GC-MS) as described (Jeong et al.2004). Briefly, the dried extract was methoximated in 15 μl methoxyamine hydrochloride/pyridine solution (20 mg/ml; Sigma-Aldrich) for 30 min at 30°C, then silyated for 90 min at 60°C after adding 30 μl N-Methyl-N-(trimethylsilyl) trifluoroacetamide (Sigma-Aldrich). Incubations were carried out in a Vortemp 56 orbital shaker (Labnet) at 600 rpm. Derivitized samples were injected (1 µl) in 25:1 split mode at an inlet temperature of 250°C. Metabolites were resolved on a DB-5MS column (30 m length, 0.25 mm diameter with DuraGuard pre-column) with a helium flow of 1 ml/min. GC (Agilent 7890A) oven temperature at injection was 80°C. Following a 1 min hold at 80°C, temperature was ramped 20°C/min to 200°C, then 10°C/min to 320°C with a 6.5 min hold at 320°C. Metabolites were detected using an Agilent 5975C MS with source and quadrupole mass filter temperature setting of 230°C and 150°C, respectively. Mass spectra were collected in scanning ion mode (*m/z* 50 and 500) in ChemStation (Agilent) and deconvoluted using AnalyzerPro (SpectralWorks). Peak retention times and spectral matches corresponding to fructose, glucose and sucrose were determined using authentic standards. Peaks areas were integrated using AnalyzerPro.

### Second defoliation experiment

The second experiment was conducted mainly to collect tissues for metabolite assays (starch, condensed tannins and salicinoid phenolic glycosides) that were not carried out for the first experiment due to sample deterioration during long-term storage. In addition, conducting the second experiment presented an opportunity to extend the defoliation treatment for further assessment of defoliation effects on growth. Only a 50% defoliation treatment was utilized, and the duration of the second experiment was increased to 16 days. Starch, condensed tannin and phenolic glycoside levels were measured in order to compare RNAi effects on growth with those on non-structural metabolic sinks. Plants were propagated and grown as before and were approximately the same size at the start of treatment as in the first experiment. Tissue collection and processing for metabolic analysis followed the same procedures as above.

### Starch

Ten mg of ball-milled leaf, bark or stem powder in 2 ml Eppendorf tubes were extracted five times using 1.5 ml ethanol:chloroform (1:1, v/v) with sonication. After two rinses with 80% ethanol, α-amylase (1000 U, Sigma A4551) dissolved in 0.5 ml buffer (0.1 M sodium acetate, pH 5.0, with 5mM CaCl_2_) was added to the extracted residue. After a 30 min incubation at 85°C with 800 rpm shaking, samples were cooled and 10 ul of sodium acetate buffer (pH 5.0) containing 5 U amyloglucosidase (Sigma A1602) was added to the digest. Samples were digested at 50°C for 48 hr with shaking (800 rpm). Digests were then centrifuged at 24,000 g and an aliquot of the supernatant was removed for GC-MS detection of glucose as described above.

### Condensed tannins

Condensed tannins (CT) were analyzed as in our previous work (Harding et al. 2005). Briefly, approximately 10 mg freeze-dried tissue powder was extracted in 600 μl of methanol for 15 min in an ultrasonic bath and centrifuged at 15,000 g for 10 min. Pigment-containing supernatant was combined with water (400 μl) and chloroform (400 μl) in a new tube, then vortexed and centrifuged to remove pigments. The depigmented pellet and supernatant were combined and dried down for CT analysis by the butanol-HCl method (Porter et al. 1985). Following incubation of the residues at 95°C for 20 min in 1 ml butanol-5% hydrochloric acid containing ferric ammonium sulfate, absorbance (A_550_) was read and quantified against aspen leaf CT standards.

### Phenolic glycosides

Approximately 10 mg ball-milled, freeze-dried tissue leaf or bark tissue was used for each assay. Powders were sonicated twice for 15 min each at 4°C in 400 μl of a master mix containing chloroform:methanol (1:1, v/v) and 700 μM D_5_-benzoic acid internal standard. Water (200 μl) was added to the sonicate which was vortexed and centrifuged to obtain a pigment-free upper phase for HPLC-TOF analysis. Samples were chromatographed using a Zorbax Eclipse XDB-C18 High Resolution 4.6×50 mm column with 1.8 micron particle size. Mobile phases were (A) 97% H_2_O/3% acetonitrile/0.1% formic acid and (B) 97% acetonitrile/3% H_2_O/0.1% formic acid. Column temperature was 30°C. Elution followed the gradient: 3%B (0-1 min); 3%B-17%B (1-3 min); 17%B (3-5 min); 17%B-60%B (5-9 min); 60%B-98%B (9-11 min). Detection was by MS-TOF (Agilent 6220) in negative mode using electro-spray ionization, capillary voltage 3500V, and fragmenter voltage set at 125V. Retention times and MS spectra were compared to those of salicortin (formic acid adduct *m/z* 470.4) and tremulacin (formic acid adduct *m/z* 573.5) isolated from bark tissue.

### Water uptake

Water uptake was determined gravimetrically by changes in pot weight over a 90-min uptake period shortly before plants were to be harvested. Water uptake was calculated on a leaf biomass basis. From previous work, specific leaf area (cm/g dry weight) did not differ between lines (Frost et al. 2012).

### Relative water content experiment

Relative water content (RWC) was determined as (fresh weight-dry weight)/(hydrated weight-dry weight). Hydrated weight was obtained after allowing leaves to equilibrate in a dark chamber with petioles immersed in water for 4 hr. Additional equilibration did not measurably increase hydrated weight. Primary internode sections were floated on water for the 4-hr hydration, then blotted dry, weighed and dried in a forced air oven at 60°C for 48 hr.

## Results

### Defoliation affected leaf photosynthesis similarly in WT and RNAi plants

Defoliation treatments were carried out to reduce source capacity relative to sink size. Post-defoliation carbohydrate, water uptake, and growth adjustments in WT and *SUT4*-RNAi lines were compared. At the time of treatment initiation, the stem was marked at the LPI-0 node corresponding to the first unfurled leaf reaching 2 cm in length (Larson and Isebrands 1971). Removal of every third leaf below the mark comprised the 25% defoliation treatment; and of every other leaf, the 50% defoliation treatment. The maximum photosynthesis rate of post-treatment emerged source leaf (LPI-6) did not change significantly eight days after 25% defoliation in any of the genotypes. Trends toward increased photosynthesis were observed in all plants by the 50% defoliation treatment, but the increases were statistically significant only in RNAi line G, nearly so in line F, and not in WT (Table 1). When data from all three lines were pooled together, photosynthesis increased significantly by ∼15% (*P* = 0.001) in the 50% defoliation treatment. Photosynthesis of LPI-6 did not change significantly following a 100% defoliation treatment in which all leaves below LPI-0 were removed, but trended lower than in undefoliated or partially defoliated plants (Table 1). Internal leaf CO_2_ concentrations trended lower in all lines as photosynthesis increased following 50% defoliation (Table 1).

**Table 1.**
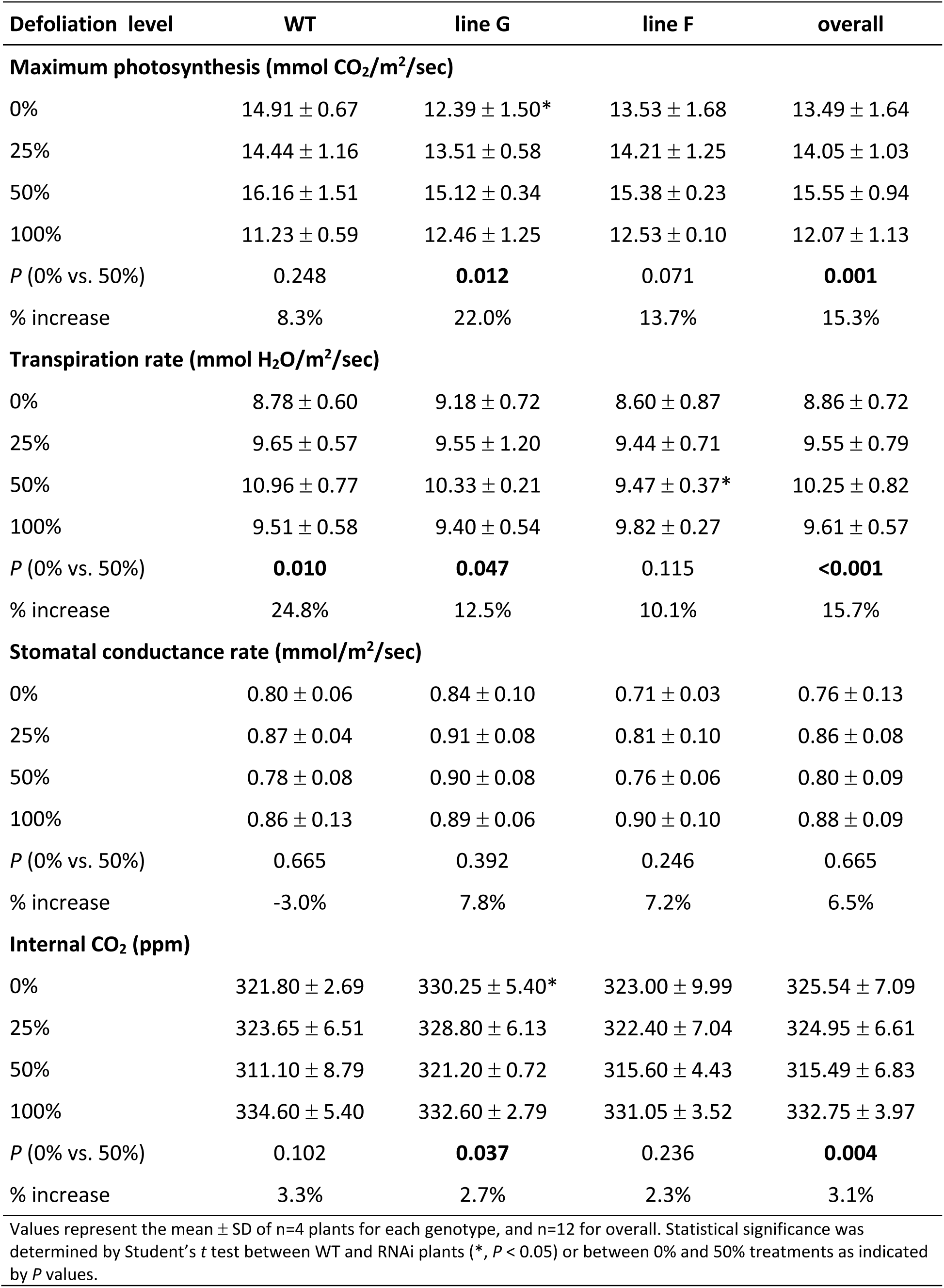
Leaf gas exchange after 8 days at various defoliation levels

| Defoliation level | WT | line G | line F | overall |
| --- | --- | --- | --- | --- |
| **Maximum photosynthesis (mmol CO<sub>2</sub>/m<sup>2</sup>/sec)** |  |  |  |  |
| 0% | 14.91 ± 0.67 | 12.39 ± 1.50* | 13.53 ± 1.68 | 13.49 ± 1.64 |
| 25% | 14.44 ± 1.16 | 13.51 ± 0.58 | 14.21 ± 1.25 | 14.05 ± 1.03 |
| 50% | 16.16 ± 1.51 | 15.12 ± 0.34 | 15.38 ± 0.23 | 15.55 ± 0.94 |
| 100% | 11.23 ± 0.59 | 12.46 ± 1.25 | 12.53 ± 0.10 | 12.07 ± 1.13 |
| *P* (0% vs. 50%) | 0.248 | **0.012** | 0.071 | **0.001** |
| % increase | 8.3% | 22.0% | 13.7% | 15.3% |
| **Transpiration rate (mmol H<sub>2</sub>O/m<sup>2</sup>/sec)** |  |  |  |  |
| 0% | 8.78 ± 0.60 | 9.18 ± 0.72 | 8.60 ± 0.87 | 8.86 ± 0.72 |
| 25% | 9.65 ± 0.57 | 9.55 ± 1.20 | 9.44 ± 0.71 | 9.55 ± 0.79 |
| 50% | 10.96 ± 0.77 | 10.33 ± 0.21 | 9.47 ± 0.37* | 10.25 ± 0.82 |
| 100% | 9.51 ± 0.58 | 9.40 ± 0.54 | 9.82 ± 0.27 | 9.61 ± 0.57 |
| *P* (0% vs. 50%) | **0.010** | **0.047** | 0.115 | **<0.001** |
| % increase | 24.8% | 12.5% | 10.1% | 15.7% |
| **Stomatal conductance rate (mmol/m<sup>2</sup>/sec)** |  |  |  |  |
| 0% | 0.80 ± 0.06 | 0.84 ± 0.10 | 0.71 ± 0.03 | 0.76 ± 0.13 |
| 25% | 0.87 ± 0.04 | 0.91 ± 0.08 | 0.81 ± 0.10 | 0.86 ± 0.08 |
| 50% | 0.78 ± 0.08 | 0.90 ± 0.08 | 0.76 ± 0.06 | 0.80 ± 0.09 |
| 100% | 0.86 ± 0.13 | 0.89 ± 0.06 | 0.90 ± 0.10 | 0.88 ± 0.09 |
| *P* (0% vs. 50%) | 0.665 | 0.392 | 0.246 | 0.665 |
| % increase | -3.0% | 7.8% | 7.2% | 6.5% |
| **Internal CO<sub>2</sub> (ppm)** |  |  |  |  |
| 0% | 321.80 ± 2.69 | 330.25 ± 5.40* | 323.00 ± 9.99 | 325.54 ± 7.09 |
| 25% | 323.65 ± 6.51 | 328.80 ± 6.13 | 322.40 ± 7.04 | 324.95 ± 6.61 |
| 50% | 311.10 ± 8.79 | 321.20 ± 0.72 | 315.60 ± 4.43 | 315.49 ± 6.83 |
| 100% | 334.60 ± 5.40 | 332.60 ± 2.79 | 331.05 ± 3.52 | 332.75 ± 3.97 |
| *P* (0% vs. 50%) | 0.102 | **0.037** | 0.236 | **0.004** |
| % increase | 3.3% | 2.7% | 2.3% | 3.1% |
| Values represent the mean ± SD of n=4 plants for each genotype, and n=12 for overall. Statistical significance was determined by Student's *t* test between WT and RNAi plants (*, *P* < 0.05) or between 0% and 50% treatments as indicated by *P* values. |  |  |  |  |

### Defoliation affected transpiration water uptake and leaf water retention differently in WT and RNAi plants

LPI-6 transpiration rates were significantly higher overall after 8 days in 50% defoliated than in non-defoliated controls in Experiment 1 (Table 1). In contrast to what we observed for photosynthesis, the transpiration increase was larger in WT than RNAi lines by roughly 2-fold (25% and 10-13%, respectively) (Table 1). The increase was significant for WT, but for only one of the RNAi lines. Stomatal conductance did not differ between genotypes or defoliation treatments (Table 1). Transpiration was not measured in Experiment 2, but water uptake normalized to leaf mass was measured in that experiment and repeated in a separate cohort of plants that were defoliated and harvested two weeks after the first cohort. In both cases, water uptake was measured just before harvest at 16 days (Figure 1). Water uptake was similar in non-defoliated WT and RNAi plants in both trials, but the increase following 50% defoliation was approximately 2-fold greater in WT than in plants of either RNAi line (Figure 1).

**Figure 1.**
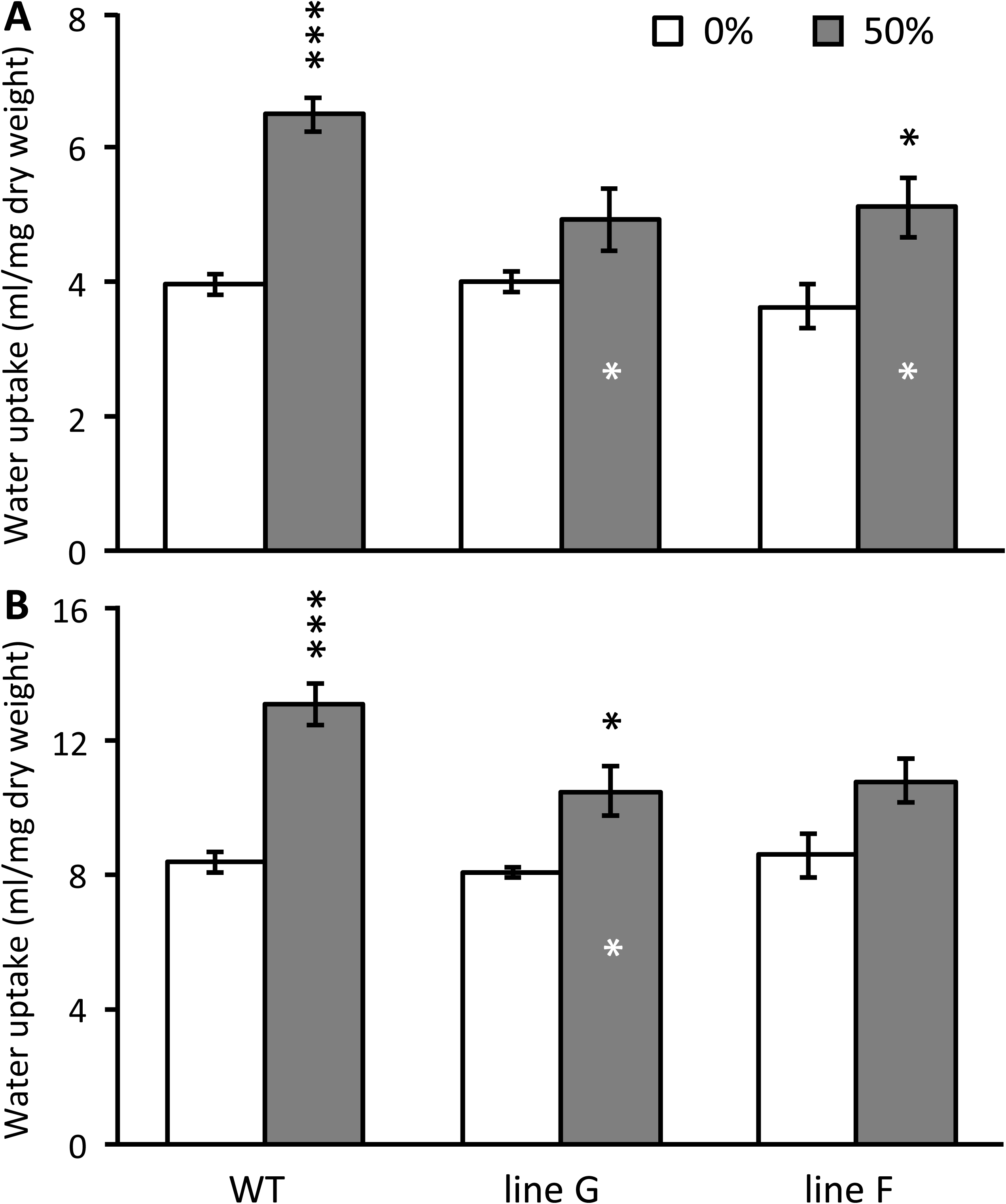
Partial defoliation increased plant water uptake on a leaf mass basis. Water uptake was measured gravimetrically over a 90-min period at mid-day (A), and for a separate cohort of plants over a 150-min period at mid-day (B). Data represent the mean and standard error of n=5 WT or n=3 RNAi plants. Asterisks over bars indicate significant defoliation effects as determined by Student’s *t* test, and asterisks inside the bar indicate significant differences between WT and RNAi lines at 50% defoliation (*** *P* < 0.001, * *P* < 0.05).

Leaf water contents were measured only in Experiment 1. Defoliation led to increased water content in new leaves that emerged and expanded after defoliation (Figure 2A). The increases were significant at all defoliation levels for the RNAi lines, but only at 100% defoliation for WT. At 100% defoliation, the % water increases were the same in all lines. Source leaves that remained on the plant after partial defoliations of 25% and 50% exhibited water content decreases in WT but not RNAi lines (Figure 2B).

**Figure 2.**
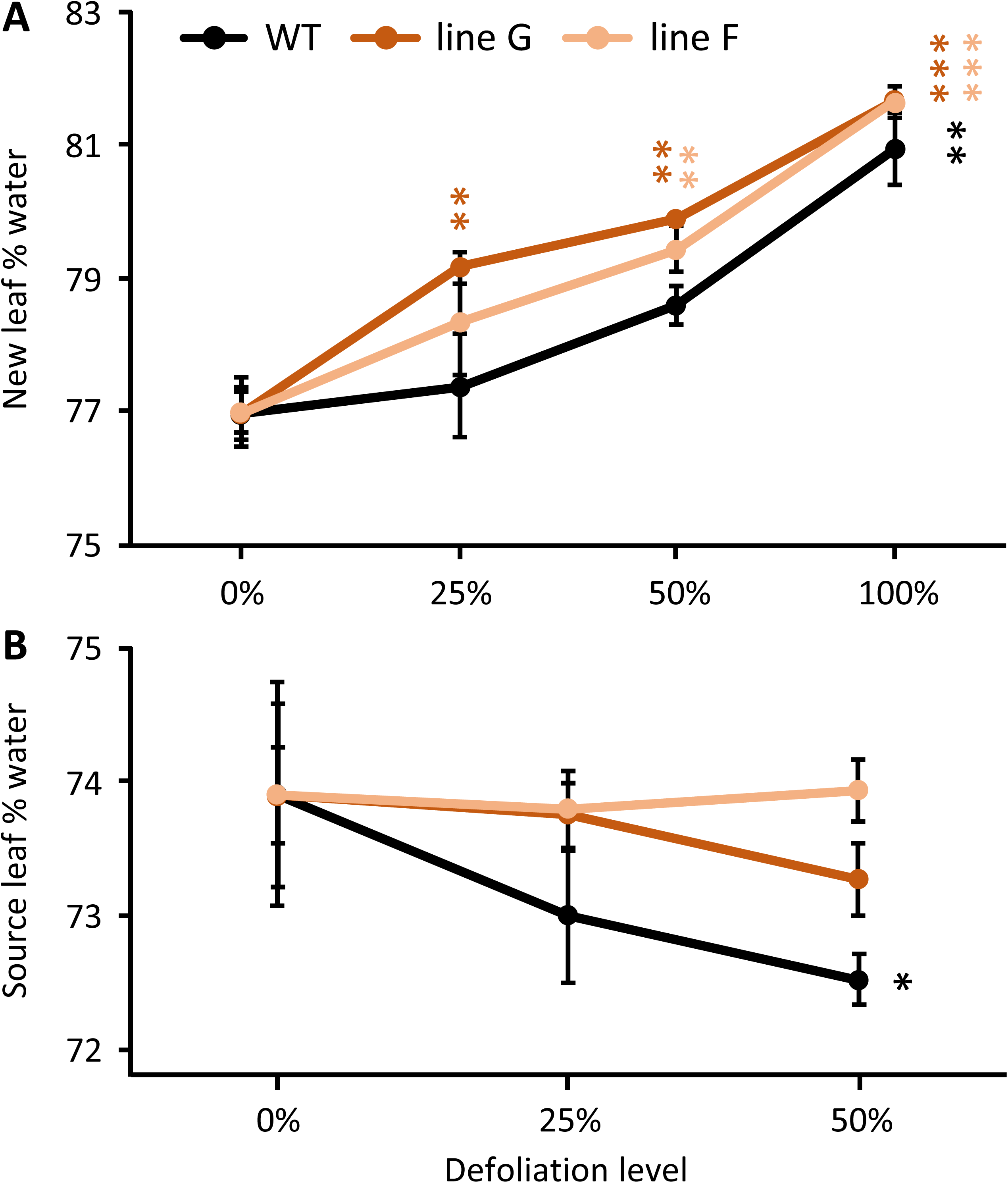
Leaf water contents responded differentially to partial defoliation in WT and RNAi plants. (A) New leaves that expanded during the defoliation treatment. (B) Source leaves that were fully expanded at the start of the defoliation treatment. The data points are connected by lines to depict the trend in water content change as defoliation levels increased for each plant line. Data represent the mean and standard error of n=4 plants. Asterisks are color coded by plant line and indicate significant defoliation effects versus non-defoliated (0%) controls as determined by Student’s *t* test (*** *P* < 0.001, ** *P* < 0.01, * *P* < 0.05).

### Defoliation affected sucrose and hexose levels similarly in WT and RNAi plants

Prior to defoliation, sucrose levels were higher in the RNAi lines as has been reported (Payyavula et al. 2011). The 8-day experiment tested three defoliation levels, 25%, 50% and 100%, and the only sustained trend across all three defoliation levels was decreasing soluble sugars (hexose and sucrose) in wood (Figure 3). The decrease was apparent in both WT and RNAi plants. Similar patterns were observed after a 16-day 50% defoliation treatment (Experiment 2), with trends toward decreased soluble sugar levels in both wood and bark (Figure S1). The apparent magnitude of the decrease after 50% defoliation (Experiment 1 and Experiment 2) or 100% defoliation (Experiment 1 only) differed little between lines (Figure 3 and Figure S1). The average sugar decreases after 8 and 16 days of partial (50%) defoliation are summarized in condensed form for all shoot organs to illustrate the broad finding that there was no clear depletion of leaf sugars by either treatment, but that depletions became more severe after 16 days for bark, and especially wood (Table 2).

**Table 2.**
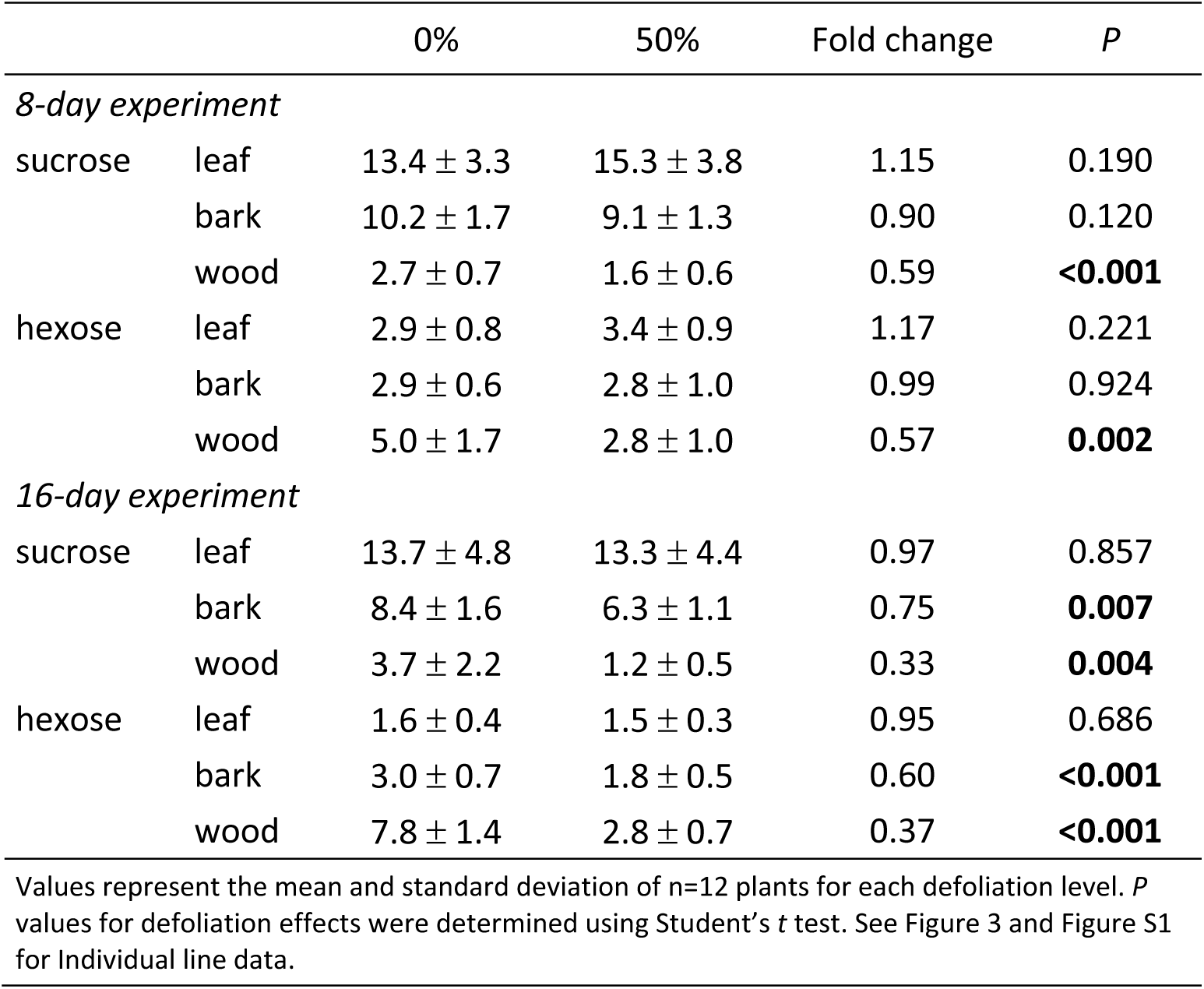
Overall changes in sucrose and hexose concentration in three shoot organs due to 8-day or 16-day defoliation treatments

**Figure 3.**
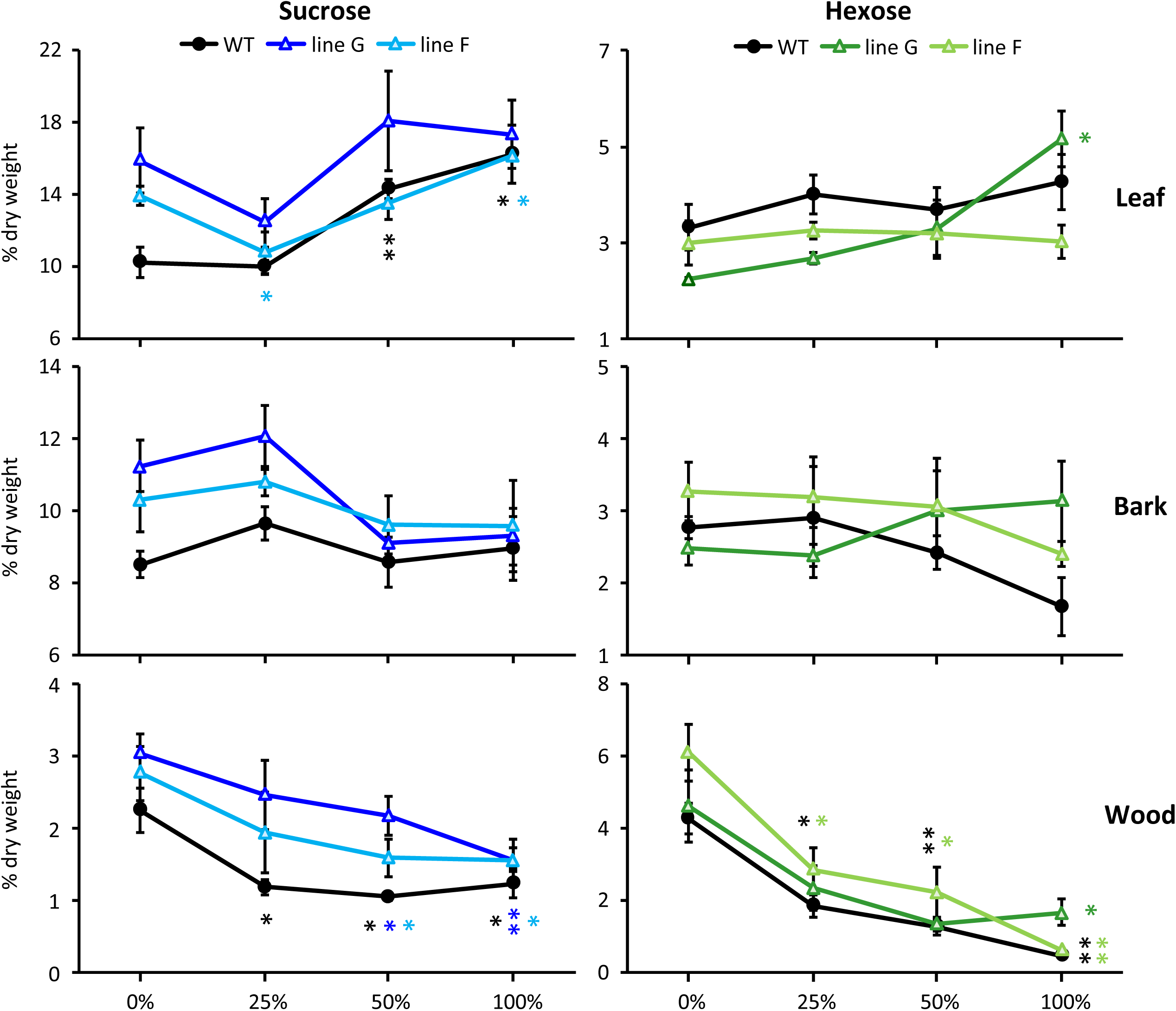
Sucrose and hexose concentration trends in leaves, bark and wood of WT and RNAi plants in Experiment 1. Data represent the mean and standard error of n=4 plants. Asterisks are color-coded by line to indicate significant defoliation effects versus non-defoliated (0%) controls as determined by Student’s *t* test (** *P* < 0.01, * *P* <0.05).

Leaf, bark and wood starch were only measured in the 16-day study (Experiment 2). By day 16, starch decreased, most clearly in bark and wood of the 50% defoliated plants (Figure 4). As with sugars, the decreases were similar in WT and RNAi lines (Figure 4).

**Figure 4.**
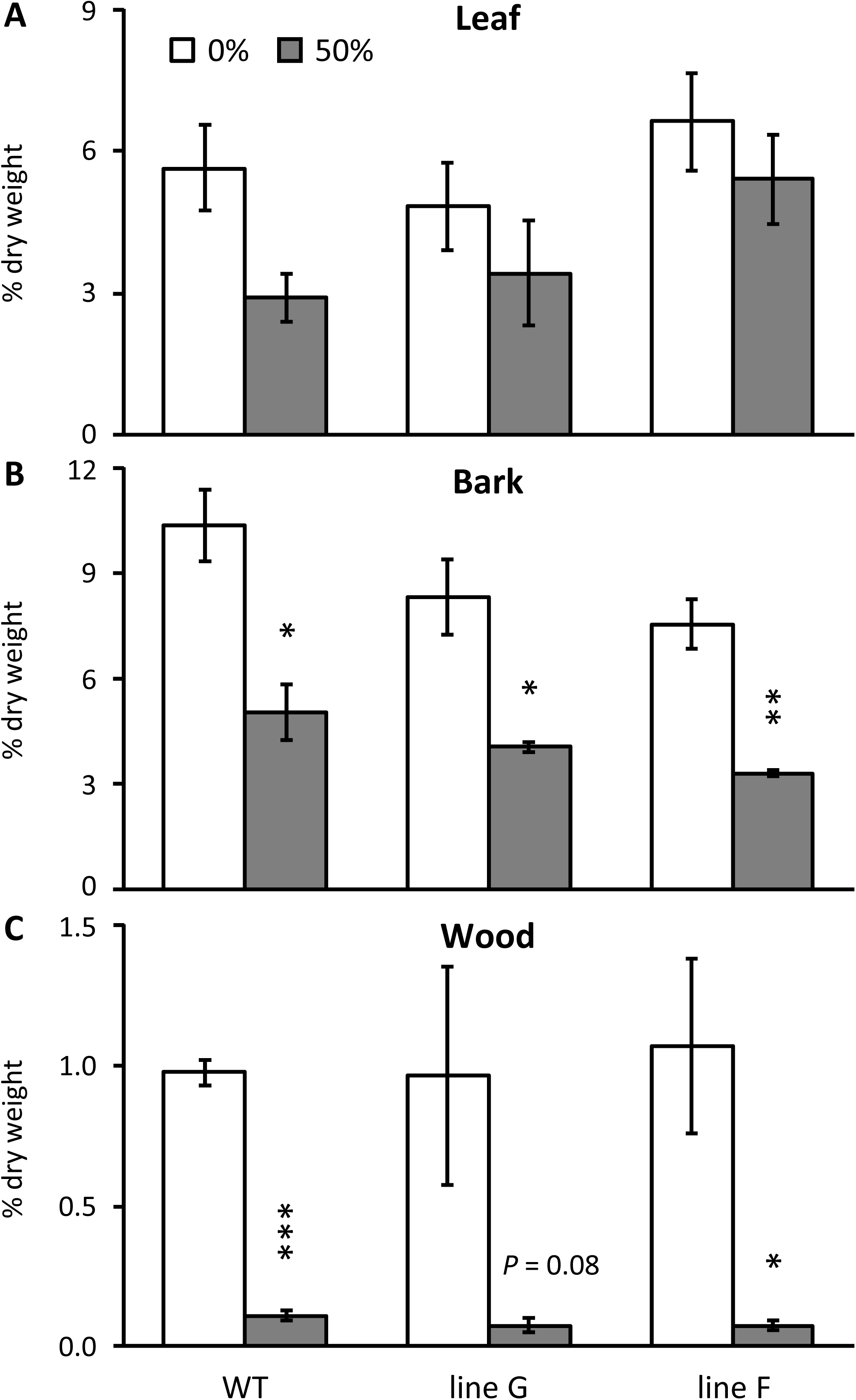
Defoliation-induced starch loss in shoot organs. Data represents the mean and standard error of n=3 plants. Asterisks indicate significant defoliation effects as determined by Student’s *t* test (*** *P* < 0.001, ** *P* < 0.01, * *P* < 0.05).

### Relative water content changed more rapidly in WT than RNAi tissues during solar warming

The transpiration and leaf water content differences between WT and RNAi plants led us to conduct a stand-alone experiment to assess whether leaf relative water content (RWC) changed the same way in WT and RNAi tissues when plants underwent gradual warming due to increasing sunlight (Table 3). Water uptake was not measured but pots were watered to saturation at the start of the 4-hr measurement period. As expected, water content decreased significantly in all lines as temperature and light intensity increased (Table 3). RWC also decreased significantly in all lines, but the decrease in upper stem internodes of RNAi was half that observed in WT plants between 7 AM and 11 AM (Table 3). While upper stem RWC did not differ statistically between genotypes at 7 AM, upper stem RWC was significantly higher in RNAi than WT plants at 11 AM. During the same 4-hr period, the RWC average in mature source leaves also decreased more in WT than RNAi plants (Table 3).

**Table 3.** Water content and relative water content changes in leaves and stem internodes during solar warming.

|  | Water content (%) |  |  |  | Relative water content (%) |  |  |  |
| --- | --- | --- | --- | --- | --- | --- | --- | --- |
|  | 7 AM | 11 AM | loss | *P* | 7 AM | 11 AM | loss | *P* |
| Internodes 5-6 |  |  |  |  |  |  |  |  |
| WT | 79.00 | 76.30 | 2.70 | **0.025** | 95.58 | 89.34 | 6.24 | **0.003** |
| RNAi | 80.80 | 78.70 | 2.10 | **0.008** | 96.47 | 93.29* | 3.18 | **0.007** |
| LPI-15 |  |  |  |  |  |  |  |  |
| WT | 72.20 | 70.30 | 1.90 | **0.037** | 98.68 | 93.40 | 5.28 | **0.001** |
| RNAi | 69.80 | 68.50 | 1.30 | **0.022** | 99.29 | 94.68 | 4.61 | **<0.001** |

Water content was determined as (fresh weight – dry weight)/fresh weight, and relative water content was (fresh weight - dry weight)/(hydrated weight-dry weight). Values represent the mean of n=5 plants. Statistical significance was determined using Student's *t* test between WT and RNAi plants (\*, *P* < 0.05) or between measurement times (indicated by *P* values).

### Secondary metabolism was not altered in RNAi compared to WT plants

Non-structural phenylpropanoid end-products, including flavonoid-derived proanthocyanidins, or condensed tannins (CT), and phenolic glycoside salicinoids typically accumulate in leaves and bark of poplars and other species of the family Salicaceae (Harding et al. 2005; Harding et al. 2014). They comprise important metabolic sinks because of their abundance, metabolic stability and impact on trophic interactions (Kleiner et al. 1999; Ruuhola and Julkunen-Tiitto 2000; Kandil et al. 2004). Average CT abundance trended lower, but not significantly, in leaves and bark of partially defoliated than of non-defoliated plants, and there was no difference between genotypes (Table 4). Salicinoids, shown as the abundance of the major phenolic glycosides salicortin and tremulacin summed, exhibited no significant differences in leaves or bark in response to partial defoliation, though there were modest small decreases in bark. In all cases, the magnitudes of the CT and salicinoid trends were similar in WT and RNAi lines.

**Table 4.** Levels of salicinoids and condensed tannins in response to defoliation treatments

|  | Salicinoids |  |  | Condensed tannins |  |  |
| --- | --- | --- | --- | --- | --- | --- |
|  | WT | Line G | Line F | WT | Line G | Line F |
| *Leaf* |  |  |  |  |  |  |
| 0% | 8.00 ± 0.88 | 7.40 ± 0.39 | 7.83 ± 0.70 | 6.22 ± 3.28 | 6.54 ± 1.25 | 6.77 ± 0.80 |
| 50% | 8.74 ± 1.22 | 7.74 ± 1.02 | 7.68 ± 0.95 | 3.44 ± 1.40 | 4.64 ± 1.81 | 5.49 ± 2.53 |
| *P* | 0.44 | 0.61 | 0.84 | 0.25 | 0.21 | 0.45 |
| *Bark* |  |  |  |  |  |  |
| 0% | 10.53 ± 0.75 | 10.97 ± 0.46 | 10.46 ± 0.57 | 2.80 ± 0.23 | 2.04 ± 0.25* | 3.22 ± 0.75 |
| 50% | 9.07 ± 0.60 | 9.25 ± 1.42 | 9.18 ± 1.33 | 2.33 ± 0.58 | 2.01 ± 0.50 | 2.18 ± 0.39 |
| *P* | 0.06 | 0.12 | 0.3 | 0.26 | 0.95 | 0.1 |

Values represent the mean±SD of n=3 plants. Statistical significance was determined by Student's *t* test between WT and RNAi plants (\*, *P* < 0.05) or between 0% and 50% treatments shown by *P* values. Salicinoids represent the sum of salicortin and tremulacin levels.

### Shoot growth was more negatively affected by partial defoliation in RNAi than WT

Stem biomass was divided by leaf biomass to obtain an index of leaf ability to sustain stem biomass production, or ‘growth’ in Experiment 2. Stem mass per unit leaf mass did not differ between WT and RNAi lines when intact plants were compared, but was greater for WT than either RNAi line when defoliated plants were compared (Figure 5).

**Figure 5.**
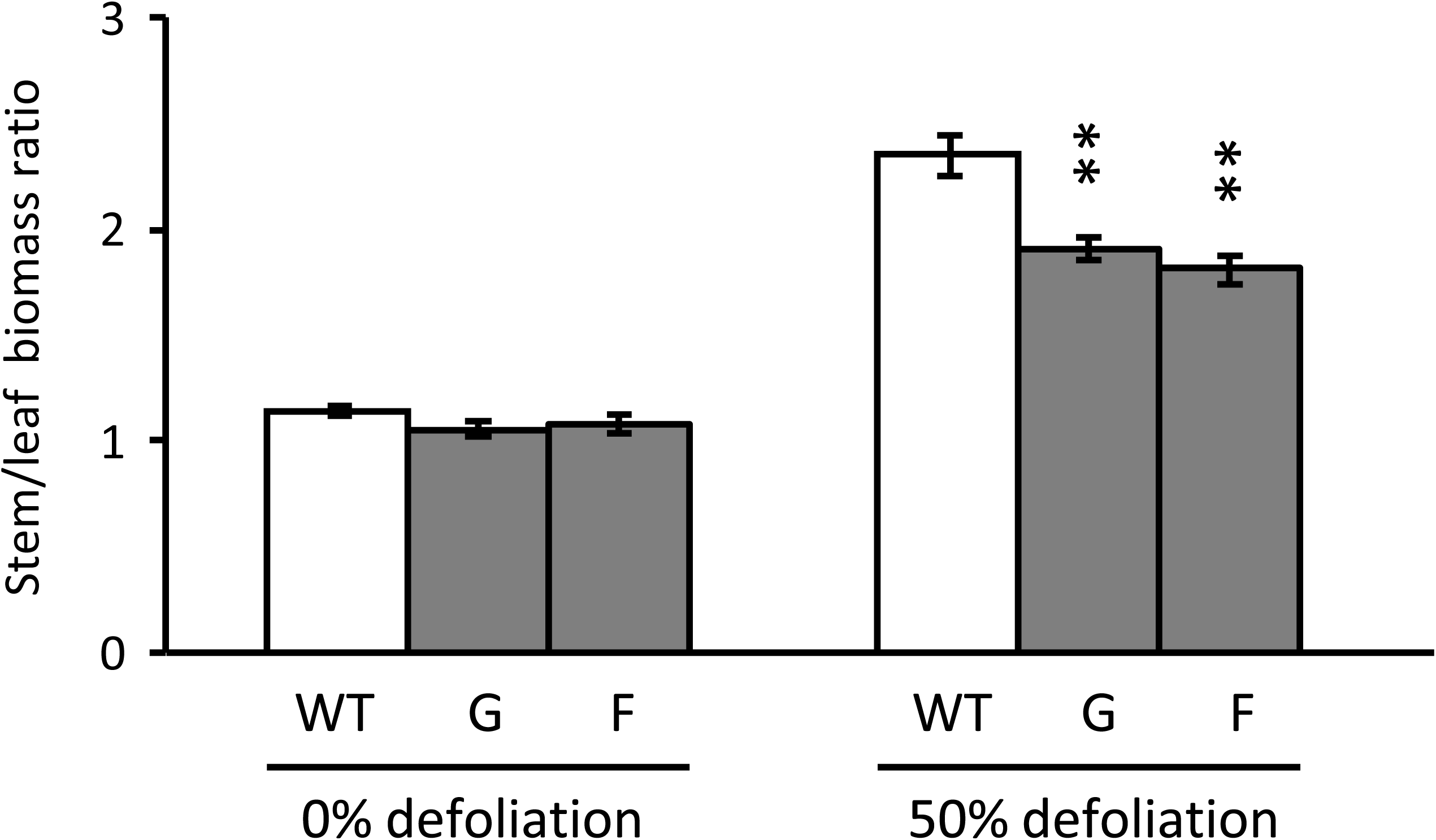
Stem growth per unit leaf mass was higher in wild type than RNAi plants following a 16-day 50% defoliation treatment in Eexperiment 2. Data represent the mean and standard error of n=5 WT or n=3 RNAi plants. Asterisks indicate significant growth differences between partially defoliated WT and RNAi plants as determined by Student’s *t* test (** *P* < 0.01).

## Discussion

The knowledge gap we sought to address centers around the physiological role of PtaSUT4, a highly conserved tonoplast sucrose transporter with orthologs in monocots, dicots and basal angiosperm taxa (Peng et al. 2014). While SUT4 mediates sucrose efflux from the vacuole into the cytosol, the physiological relevance of that efflux in tree species such as *Populus* is being revealed only gradually (Payyavula et al. 2011; Frost et al. 2012; Xue et al. 2016). Those studies explored the effects of SUT4-RNAi on overall plant growth, sugar level and gene expression under varied conditions of water or nitrogen deficit. The defoliation treatments in this study were expected to increase transpiration and thus water flux through the remaining leaves as has been reported elsewhere for *Populus* (Liu et al. 2014), thereby testing SUT4 function in a novel context with new physiological implications. In short, leaf water flux increased, but less robustly in RNAi than WT plants (Table 1 and Figure 1). Hydraulic conductivity through roots can also be decreased by partial defoliation in *Populus* (Liu et al. 2014). Although root conductivity was not measured in our study, the greater increases we observed in RNAi leaf water content following defoliation (Figure 2) are not consistent with the idea of a more limiting root hydraulic conductivity in the RNAi lines. In contrast to the differentials in water uptake, sink carbohydrate levels decreased similarly after defoliation in all genotypes (Table 2, Figures 3-4, and Figure S1). Photosynthesis was only measured in Experiment 1, and it increased at least as strongly in RNAi as in WT plants after partial defoliation (Table 1). In light of the differences in water uptake, we concluded that the greater growth reductions in RNAi than WT plants after defoliation (Figure 5) were due more to water uptake than to carbohydrate availability.

Because leaf water content increased more in RNAi plants, and stomatal conductance did not vary between lines or defoliation levels, we suggest that less robust compensatory uptake of water by the RNAi plants (Table 1 and Figure 1) reflected a weaker capacity to increase leaf hydraulic conductivity, the ability of water to move through the leaf toward the stomata. In a previous study, we reported that plant water uptake was slower in RNAi plants even in the absence of a defoliation treatment (Frost et al. 2012). However, the water uptake differences in that study were observed in a context of soil drying. The findings from both studies are consistent with a slowed or limited ability of RNAi leaves to make hydraulic conductivity adjustments.

Partial defoliation and silvicultural treatments such as thinning have long been known to result in increased photosynthesis, transpiration and productivity on a leaf area or plant basis (von Caemmerer and Farquhar 1984; Breda et al. 1995). Increased transpiration under non-stress conditions is supported by increased hydraulic conductivity which minimizes the lowering of leaf water potentials in a way that might be detrimental to photosynthesis (Brodribb and Holbrook 2006; Blackman et al. 2009; Liu et al. 2014; Simonin et al. 2015). Decreased leaf water potential has been reported in the case of well-watered, partially defoliated *Eucalyptus* when hydraulic conductivity did not increase (Eyles et al. 2013). A number of studies suggest that dynamic adjustments in hydraulic conductivity depend on the expression of aquaporins (Cochard et al. 2007; Pou et al. 2013; Liu et al. 2014). Aquaporins form pores in membranes that allow water to move intracellularly or across membranes and through tissues toward locations of lower water potential (Kapilan et al. 2018). The degree to which such adjustments can alter hydraulic conductivity will also depend on anatomical traits that differ between species and dictate the mass movement of water within tissues, or a plant’s vulnerability to embolism and hydraulic failure (Zwieniecki et al. 2007; Blackman et al. 2009).

In our study, leaves that emerged and expanded before (Figure 2A) as opposed to after (Figure 2B) defoliation certainly differed with respect to their water retention responses. However, there was a tendency toward higher water retention by *SUT4*-RNAi leaves than WT leaves, regardless of their developmental stage (Figure 2A versus 2B) at the time of defoliation. This points toward a dependence of at least part of the RNAi effect on a subcellular dynamic that did not necessarily depend on developmental changes in vascular and leaf surface characteristics. In fact, decreases in RWC developed more rapidly in WT than RNAi tissues in response to gradual warming and increasing light intensity during a 4-hr period (Table 3). RWC correlated similarly with leaf water potential measured by a pressure bomb for both WT and RNAi leaves (Figure S2). At this point therefore, we interpret the findings to be consistent with the idea that reduced inter-compartmental sucrose trafficking has consequences for the dynamic control of osmotic gradients within the cell. Although we can only offer this as a suggestion, it is possible that the export of vacuolar sucrose into the cytosol is coupled with increased cytosolic invertase activity. This in turn would be expected to affect osmotic gradients and the expression or activity of aquaporins that condition hydraulic conductivity and ultimately transpiration (Chaumont and Tyerman 2014; Maurel et al. 2015).

Following this line of reasoning, we speculate that sucrose export from the vacuole to the cytosol in parenchyma or mesophyll cells that are adjacent to minor vein xylem vessels in the leaf could promote withdrawal of water from the vessels into the extraxylary symplasm. Aquaporins could then facilitate the cross-membrane dispersal of water via extraxylary apoplastic and symplastic spaces throughout the leaf, leading to enhanced transpiration. The idea that specifically localized tonoplast aquaporins can contribute to leaf hydraulic conductivity in this fashion has been proposed (Laur and Hacke 2014). Other studies report that additional aquaporins are likely involved with the regulation of leaf hydraulic conductivity and the extraxylary movement of water (Pou et al. 2013). We have reported previously that RNAi-silencing of *PtaSUT4* indeed influences the expression of several aquaporins during a drought response (Xue et al. 2016). Under circumstances of reduced *SUT4* expression such as in RNAi plants or water-limited WT plants (Frost et al. 2012), defoliation-induced changes in aquaporin expression or activity could result in the vacuolar entrapment of water arriving from the apoplasm. The present work affirms that source reduction due to partial defoliation stimulates transpiration, but that SUT4-mediated subcellular sucrose distribution could affect the partitioning of water during its transit through the leaf.

Stronger water retention in both newly expanded and older source leaves of RNAi plants following defoliation is consistent with increased vacuolar sequestration of water entering the leaf symplasm during the transpiration increase. Greater sequestration would have the effect of competing with transpiration in the RNAi plants. We note that the magnitude of such competition could vary with the expression level of *PtaSUT4* in WT plants. For example, partial defoliation of drought-adapted poplar plants where *PtaSUT4* expression is low (Frost et al. 2012) might result in a smaller compensatory transpiration increase than in moisture replete plants. How SUT4 and aquaporins coordinate at a micro-anatomical level to condition leaf hydraulic properties in the ways suggested here is clearly beyond the scope of this work and remains to be investigated.

## Supplemental Data

**Supplemental Figure S1.**
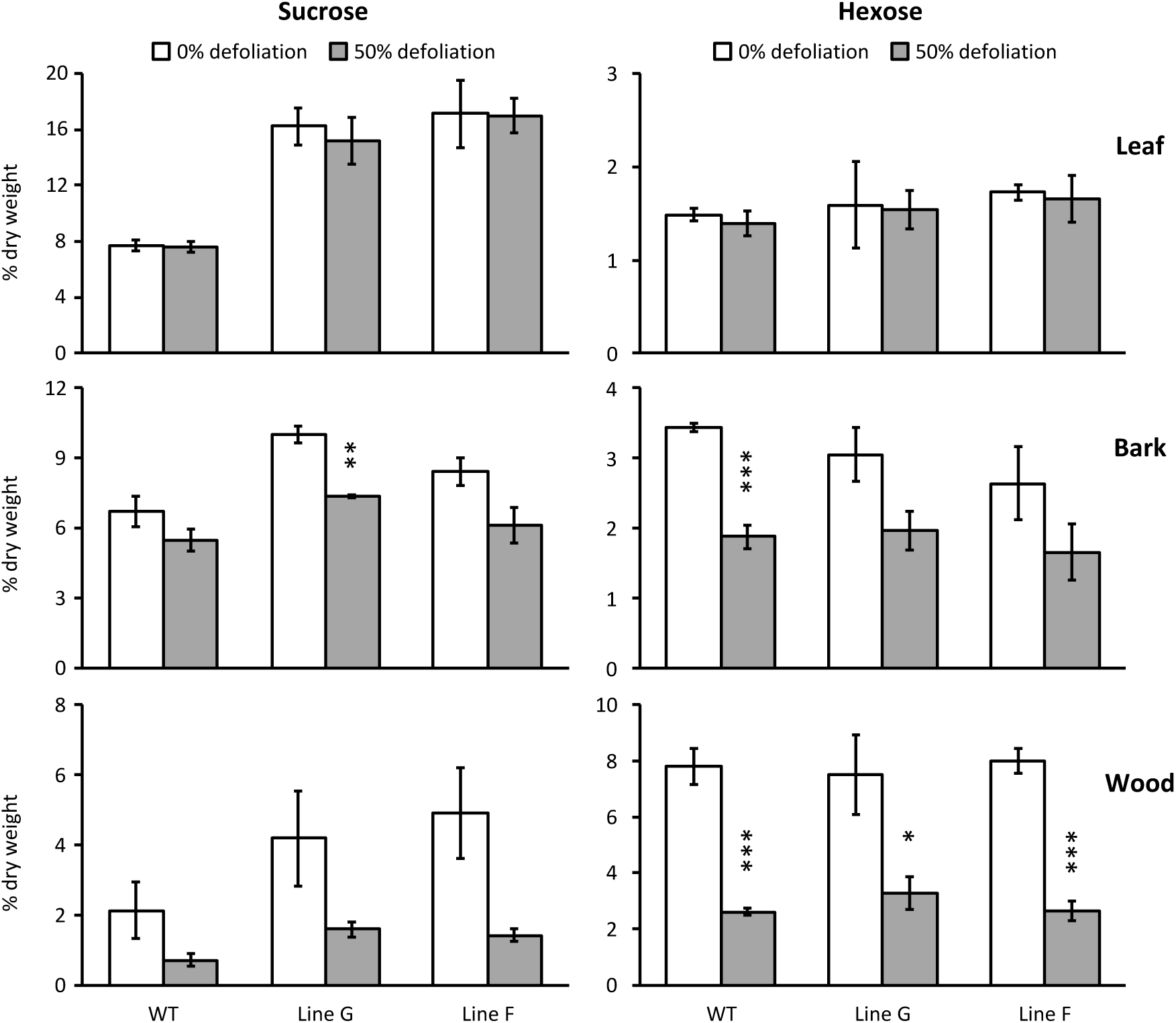
Sucrose and hexose concentration trends in leaves, bark and wood of WT and RNAi plants in Experiment 2. Data represent the mean and standard error of n=3 plants. Asterisks indicate significant defoliation effects as determined by Student’s *t* test (*** *P* < 0.001, ** *P* < 0.01, * *P* < 0.05).

**Supplemental Figure S2.**
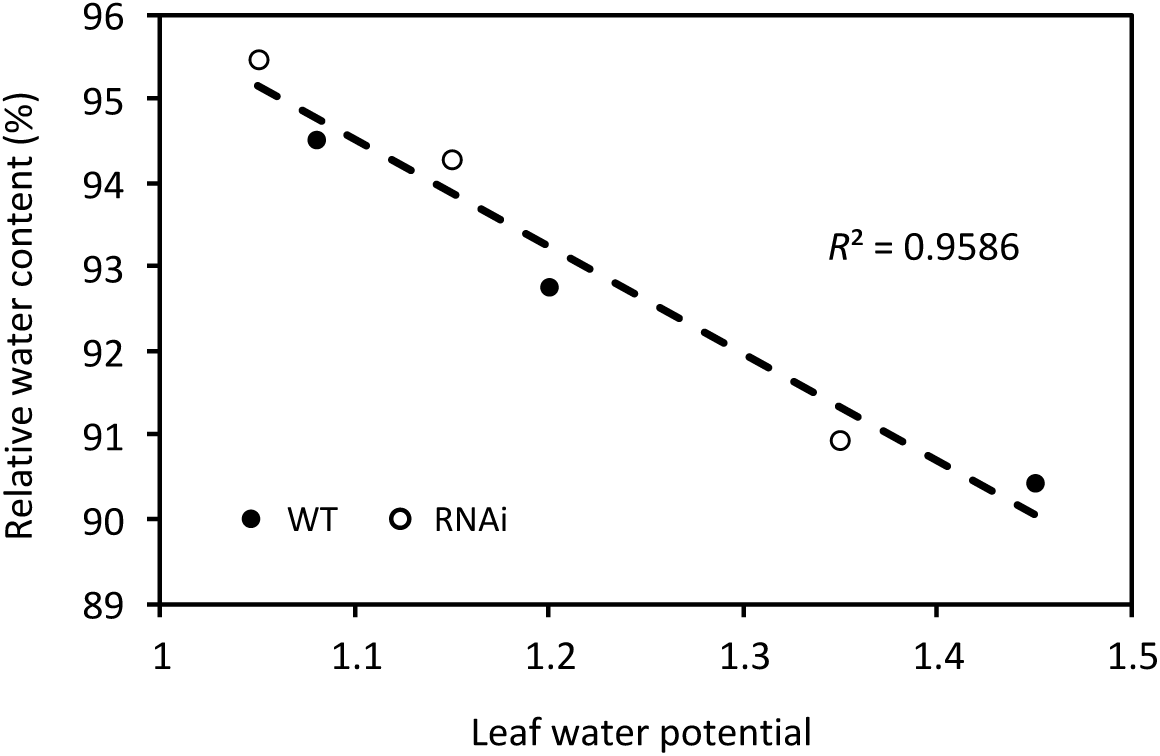
Relative water content correlated with leaf water potential as determined using a pressure bomb device. The correlation was very similar for WT and RNAi plants.

## Acknowledgements

The authors thank Stephen Pettis and Suzzanne Tate for greenhouse plant care, and Batbayar Nyamdari for GC-MS and LC-MS analysis. This research was supported by the Office of Biological and Environmental Research within the Department of Energy (grant no. DE-SC0005140) to SAH and CJT.

## Author Contributions

SAH and CJT conceived the study, CJF and SAH designed and performed the research, SAH analyzed the data and wrote the paper with contributions from CJF and CJT.

